# Culture and genomic analysis of *Symbiopectobacterium purcellii*, gen.nov. sp. nov., isolated from the leafhopper *Empoasca decipiens*

**DOI:** 10.1101/2022.01.15.476461

**Authors:** Pol Nadal-Jimenez, Stefanos Siozios, Nigel Halliday, Miguel Cámara, Gregory D.D. Hurst

**Affiliations:** Institute of Infection, Veterinary and Ecological Sciences, University of Liverpool, Liverpool, UK; The National Biofilms Innovation Centre, School of Life Sciences, University of Nottingham Biodiscovery Institute, University of Nottingham, UK

**Author notes:** **Correspondence:** Pol Nadal-Jimenez, University of Liverpool.

**Keywords:** *Symbiopectobacterium*, *Empoasca*, symbiosis, leafhopper, quorum sensing

## Abstract

Bacterial endosymbionts are found in multiple arthropod species, where they play crucial roles as nutritional symbionts, defensive symbionts or reproductive parasites. Recent work has highlighted a new clade of heritable microbes within the gammaproteobacteria that enter into both obligate and facultative symbioses, with an obligately required unculturable symbiont recently given the name *Cand.* Symbiopectobacterium. In this study, we describe a culturable rod shaped non-flagellated bacterial symbiont from this clade isolated from the leafhopper *Empoasca decipiens*. The symbiont is related to the transovarially-transmitted ‘BEV’ bacterium that was first isolated from the leafhopper *Euscelidius variegatus* by Alexander Purcell, and we therefore name the symbiont *Symbiopectobacterium purcellii* sp. nov. gen. nov. We further report the closed genome sequence for *S. purcellii*. The genome is atypical for a heritable microbe, being large in size, without profound AT bias and with little evidence of pseudogenization. The genome is predicted to encode Type II, III and VI secretion systems and associated effectors and a non-ribosomal peptide synthase array likely to produce bioactive small molecules. Predicted metabolism is more complete than for other symbionts in the *Symbiopectobacterium* clade, and the microbe is predicted to synthesize a range of B vitamins. However, Biolog plate analysis indicate metabolism is depauperate compared to the sister clade, represented by *Pectobacterium carotovorum*. A quorum-sensing pathway related to that of *Pectobacterium* spp. (containing an overlapping *expI-expR1* pair in opposite directions and a “solo” *expR2*) is evidenced, and LC-MS/MS analysis reveals the presence of 3-hydroxy-C10-HSL as the sole *N*-acylhomoserine lactone (AHL) in our strain. This AHL profile is profoundly divergent from that of other *Erwinia* and *Pectobacterium* spp., which produce mostly 3-oxo-C6- and 3-oxo-C8-HSL and could aid group identification. Thus, this microbe denotes one that has lost certain pathways associated with a saprophytic lifestyle but represents an important baseline against which to compare other members of the genus *Symbiopectobacterium* that show more profound integration into host biology.

## INTRODUCTION

It is now understood that microbes influence multiple aspects of animal biology (1). Symbiont contributions extend from involvement in the process of digestion in the gut, through anabolic activities and the supply of vitamins and amino acids, to protection against natural enemies and defence against prey/hosts (2). Conversely, other symbiotic microbes are pathogenic or parasitic, and many symbioses combine both parasitic and beneficial aspects. Levels of symbiont integration vary between symbioses (3). On the host axis, they vary from facultative relationships where the host does not require a particular symbiont, to obligate where the individual dies or becomes sterile in the absence of symbiosis. Likewise, symbionts vary in the degree to which they rely on a host – some only replicating within hosts with others having environmental replication. The process of symbiosis formation also varies – from arising within the host lifecycle through acquisition by the host or infection by the microbe, to being present through the host lifecycle, with symbiont transfer/transmission from parent to offspring.

Whilst arthropod-microbe symbioses are diverse in terms of the microbial partners, particular microbial taxa have established symbiosis with a number of host species, commonly establishing in new host species through a host switch event. Well-known ‘heritable symbionts’ found over a broad range of arthropods include *Wolbachia*, *Rickettsia*, *Spiroplasma*, *Cardinium* and *Arsenophonus* (4). The interactions found in these symbioses include obligate and facultative associations, and ones which are beneficial, parasitic or have a combination of features.

Recent research has added a new clade of insect symbionts, *Cand.* Symbiopectobacterium, to ‘the big five’ (5). The first member of this clade to be described was the BEV strain – an acronym for bacterium from *Euscelidius variegatus*, a planthopper host species. This strain was cultured (6), but never formally named. The symbiosis was characterized as one with vertical transmission, where the host’s reproduction was negatively impacted by the microbe. In addition, there was also transmission to other insects on the plant – thus establishing the symbiosis as a pathogenic one maintained through mixed modes of transmission (7). Experiments also suggested the symbiont facilitated the transmission of phytoplasma from its bug host to plant (8). Later, the pest species *Cimex lectularius* (common bedbug) was observed to carry a heritable symbiont related to BEV (9). This symbiont has not been established in cell-free culture, and symbiosis is facultative from the host perspective: the bedbug does not require the symbiont. Following this, a third hemipteran – the bulrush bug *Chilacis* was observed to carry a related vertically transmitted symbiont housed in a gut mycetome, in what appears to be an obligate association, in which the host requires the symbiont (10). More recently, symbioses involving members of this clade have extended beyond Hemiptera hosts to nematodes, with Martinson et al (5) characterizing symbionts related to BEV as obligate partners of *Howardula* nematodes. They named this microbe *Cand.* Symbiopectobacterium, reflecting its symbiotic lifestyle and its sister relationship to the well-characterized genus *Pectobacterium*.

*Cand.* Symbiopectobacterium has thus emerged as a potentially widespread and significant symbiotic associate of invertebrates. The original culturable BEV isolate, on which the genus could be formally described, was lost and genomic information for this strain is partial (11). Recovering a model culturable member of the genus is important, as it allows formal description of the microbe, completion of a closed genome sequence against which reductive evolution in symbiosis can be measured and presents a system in which gene function may be investigated. In this paper, we report the isolation to pure culture of a member of this clade from the planthopper *Empoasca decipiens*. We further present and analyse the complete genome sequence of this microbe, assess its growth requirements compared to *Pectobacterium carotovorum* and analyse its quorum sensing-signalling system.

## MATERIALS AND METHODS

### Symbiont isolation, morphology *in vitro* and identification through 16S rRNA sequence

Initial Cicadellidae samples with light green coloration were collected in Prince’s Park, Liverpool in April 2018, scooping different plants with an insect net at a maximum height of 2m. Fresh specimens were transported alive to the lab and sacrificed by freezing at −20°C for 15 min. The insect specimens were surface sterilized by immersion in 70% ethanol and washed with sterile water to remove the remaining alcohol. Insect legs were excised with a sterile surgical blade and stored at −20°C for *post hoc* host species determination through DNA barcoding.

The remainder of the insect body was mechanically crushed and resuspended in 100 μl of sterile water. An aliquot of 10 μl was plated on brain heart infusion (BHI, Oxoid, UK) agar and grown at 30°C for 6 d to allow the appearance of slow-growing bacterial colonies. Morphology was examined through Gram staining and scanning electron microscopy of overnight culture. To identify the bacterial species, we performed colony PCR of the 16S rRNA gene colonies emerging on the agar plates with primers 27F (AGAGTTTGATCMTGGCTCAG) (12) and 1492R(I) (GGTTACCTTGTTACGACTT) (13), and sequenced by Eurofins genomics, Germany. Sequences were manually curated and phylogenetic analysis performed based on the 16S rRNA gene sequences which included a large assemblage of members from the *Cand*. Symbiopectobacterium clade. To this end, 16S rRNA sequences were aligned using the SSU-ALIGN software (14). A Bayesian phylogeny was estimated with MrBayes v3.2.6 (15) by sampling across the GTR model space (nst=mixed, rates=gamma). Two independent runs were performed for 5,000,000 generations and sub-sampling every 500 generations using four Markov chains. The first 25% of the samples were discarded as burn-in.

The leafhopper host was identified through DNA barcoding using the COI sequence. To this end, insect legs were mechanically crushed and resuspended in 50 μl of sterile water and the genomic DNA (gDNA) was extracted using a Quick-DNA Universal kit (Zymo research, USA). 2 μl of the gDNA were added to a GoTaq^®^ Green Master Mix (Promega, USA) and used to amplify part of the mitochondrial cytochrome oxidase 1 (CO1) with primers C1-J-1718 (GGAGGATTTGGAAATTGATTAGTTCC) and C1-N-2191 (CCCGGTAAAATTAAAATATAAACTTC) (16). The PCR program consisted of an initial denaturation step at 95°C for 5 min, followed by 30 cycles of DNA denaturation at 94°C for 15s, primer annealing at 55°C for 45 seconds, and primer extension at 72°C for 1 min. A final extension was carried out at 72°C for 5 min. A few microliters of each PCR product were run on an agarose gel to assess the success of the PCR reaction and the remains cleaned through an Isolate II PCR and Gel kit (Bioline, USA) and sent for sequencing with primer C1-N-2191. Identity was checked through analysis against the Barcode of Life Database, BOLD.

### In vitro Growth requirements

BIOLOG GEN III plates (Cat. No. 1030) were used to ascertain the physiological and biochemical characteristics of *S. purcellii* SyEd1 *in vitro*, and these were conducted alongside *Pectobacterium carotovorum* subsp. *carotovorum* LMG 02404^T^ for comparison. Within this, we also performed the assay in the presence/absence of 0.4 % polygalacturonic acid PGA (Sigma, P3850), which is commonly used to induce the expression of plant cell wall-degrading enzymes (for preparation of PGA, see (17). For the BIOLOG GEN III assays, we used IF-A inoculating fluid (Biolog, Cat. No. 72401) with or without PGA supplementation to a final concentration of 0.4% PGA. Both bacterial species were grown overnight, diluted to an OD_600_ = 0.4 and 50 μl of this aliquot were added to a tube containing IF-A fluid. The aliquot in the IF-A tube was homogeneously mixed using a vortex and 100 μl of this suspension was added to each of the 96 wells of the BIOLOG GEN III plate. The plate was subsequently incubated at 30°C without shaking.

### Potato infection assays

*Pectobacterium* spp. are well-known plant pathogens causing soft-rot disease in several plants including potatoes, carrots and cabbages. This damage is caused by a series of secreted enzymes (cellulases, proteases, pectate lyases (Pel), pectin lyases, and polygalacturonases) commonly referred to as plant cell wall-degrading enzymes (PCWDEs). The presence of 15 putative PCWDEs and two copies of the KdgR regulator (associated to their expression) in *Symbiopectobacterium purcellii* led us to assess whether this symbiont retains the plant pathogenic activity of its sister clade, *Pectobacterium.* To this aim, virulence was tested in potatoes, using a method previously described in Nadal-Jimenez et al (18) with minor modifications. Briefly, potatoes were bought at local stores, washed with tap water, dried and surface sterilized with 70% ethanol. Slices about 0.5 cm thick were placed in sterile Petri dishes. Overnight cultures of *S. purcellii* SyEd1 and *Pectobacterium carotovorum* LMG 02404^T^ were diluted to an OD600=0.4, and 20 μl were placed at the centre of the potato slice. The same amount of sterile BHI medium was added to the negative controls. The plates were sealed with parafilm to avoid moisture loss and incubated at 25°C in dark conditions. Tissue maceration was assessed visually 24, 48 and 72 h after incubation.

### Symbiont Genome sequencing, assembly and annotation

The genome of the symbiont was completed using a combination of short (Illumina) and long (nanopore) reads by MicrobesNG (Birmingham, UK) using their enhanced genome service. Briefly, Illumina sequencing was performed using the Nextera XT library prep protocol on a HiSeq platform (Illumina, San Diego, CA, USA) using a 250bp paired end protocol. Reads were adapter trimmed using Trimmomatic 0.30, with a sliding window quality cutoff of Q15 (19). Long read genomic DNA libraries are prepared with Oxford Nanopore SQK-RBK004 kit (ONT, UK) using 400-500ng of HMW DNA and sequenced in a FLO-MIN106 (R.9.4.1) flow cell in a GridION (ONT, UK). Hybrid genome assembly of both short and long reads was performed using Unicycler version 0.4.0 under the normal mode (20). The final assembly was manually inspected for potential misassemblies by mapping the raw reads back to it. Genome annotation was performed with the NCBI Prokaryotic Genome Annotation Pipeline (21). Metabolic and functional assessment of the symbiont genome was conducted using the Kyoto Encyclopaedia of Genes and Genomes (KEGG) database (22). Identification of secondary metabolite biosynthesis gene clusters was performed using the antiSMASH server (23). Finally, prophage regions were predicted using the PHAge Search Tool Enhanced Release (PHASTER) web server (24).

### Phylogenomic analysis

The phylogenetic position of the symbiont was assessed based on the concatenated analysis of 527 single copy core proteins identified among 56 publicly available genomes. These include members of the closely related genera *Pectobacterium*, *Brenneria*, *Dickeya*, *Lonsdalea*, *Sodalis* and the recently characterized *Cand*. Symbiopectobacterium (5). Single copy orthologue protein sequences were identified using OrthoFinder v2.3.11 (25). A maximum likelihood phylogeny was inferred with IQ-TREE 2.0.3 [25] using the JTT+F+R3 substitution model selected using ModelFinder according to the Bayesian information criterion [26]. Clade support was assessed based on 1000 ultrafast bootstrap replicates [27].

### Analysis of *N-*acyl homoserine lactone synthesis

*S. purcellii* SyEd1 cultures were grown in 5 ml of BHI medium at 30°C for 24h and 200 rpm. After incubation the cultures were centrifuged and the supernatant collected and filtered through a 0.2 mm filter (SLGP033RS, Millipore). 500 μl of acidified ethyl acetate was added to 1 ml of supernatant sample and the mixture was vortexed for 1-2 min. Subsequently. The mixture was centrifuged for 1 min to allow the formation of a clear interface between the aqueous and organic layer. The organic (upper) layer was transferred using a pipette (without disturbing the aqueous layer) to a new 2ml Eppendorf. The extraction process was repeated twice more, combining the extracts for each sample into one of approximately 1.5 ml extract. Upon completion, the samples were dried under vacuum in a centrifugal evaporator.

Dried extract samples were reconstituted in 50 μl of methanol (MeOH) prior to analysis. LC-MS/MS analysis of 5 μl sample injections were conducted using a Qtrap 6500 hybrid triple-quadrupole linear ion trap mass spectrometer in tandem with an Exion LC system (Sciex). The overall method was a modification of that described by Ortori et al (26). Chromatography was achieved using a Phenomenex Gemini C18 column (3.0 um, 50 × 3.0 mm) with a constant flow rate of 450 μl/min of mobile phase A (0.1 % (v/v) formic acid) and mobile phase B (0.1 % (v/v) formic acid in methanol). The LC gradient began at 10% B for 1.0 min, increased linearly to 50% B over 0.5 min, then to 99% B over 4.0 min. The composition remained at 99% B for 1.5 min, decreased to 10% B over 0.1 min, and stayed at this composition for 2.9 min. Analyte detection was conducted with the MS operating in MRM (multiple reaction monitoring) mode, screening the LC eluent for specific AHLs (unsubstituted, 3-oxo and 3-OH AHLs with even chain lengths from C4-C14).

### Prevalence of symbiont in *Empoasca decipiens* leafhoppers

In order to assess the prevalence of this bacterium in *E. decipiens*, we performed a PCR screening on various specimens. Additional insect collections were completed at the same location in August 2019 and tested for SyEd1 by PCR assay. Using the full-genome sequence of our cultured strain, we developed two set of specific PCR primers to amplify part of the DNA gyrase subunit (*gyrB*) gene of this bacterium: BEV_gyrB_F1 (CCGTGGTGTCGGTGAAAGTA) + BEV_gyrB_R1 (TGGTCTTCTGTCAGCGTGTC) and BEV_gyrB_F2 (CTCGTGAAATGACACGACGC) + BEV_gyrB_R2 (CAGCAGTTCCACTTGTTCGC). The gDNA was extracted in the same manner as for the leg samples and used as a template for the PCR reactions.

## RESULTS

### Symbiont isolation and identification

The bacterium grows under standard aerobic conditions in Brain heart infusion (BHI) medium (CM1032, Oxoid), forming circular white colonies approx. 2-3 mm in diameter on BHI agar, and cultures emitted a pronounced plant-like odour. The bacterium is Gram negative, and SEM revealed it to be a non-flagellated rod shape, of length 1-1.5 mm (Figure 1). The bacterium will also grow in LB (Miller) (110285, Millipore/Merck KGaA) although at a slower rate, and growth is inhibited by light.

**Figure 1:**
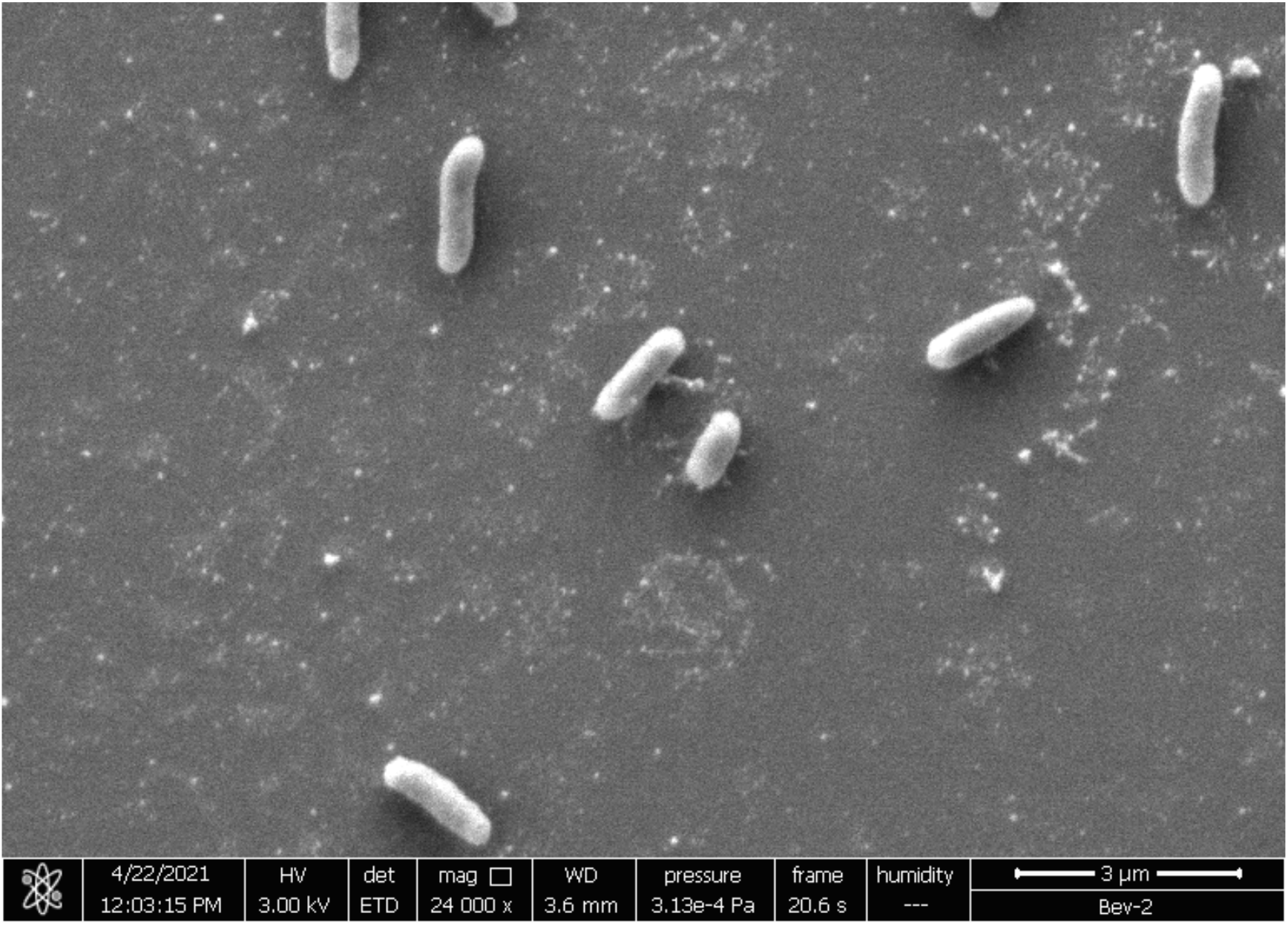
SEM of *S. purcellii* SyEd1.

Phylogenetic analysis based on the 16S rRNA gene (Accession number OK044380) placed the isolated microbe well within the recently characterized clade Cand*. Symbiopectobacterium* (Figure 2), a group of microbes commonly associated with arthropods and nematodes. Sequence of the CO-1 amplicons from the insect host revealed the leafhopper host to be *Empoasca decipiens* (Hemiptera, Cicadellidae), a common species of leafhopper in Europe. *Empoasca decipiens* has been implicated in the transmission of various plant pathogens (27), and is considered a pest in various crops (28).

**Figure 2:**
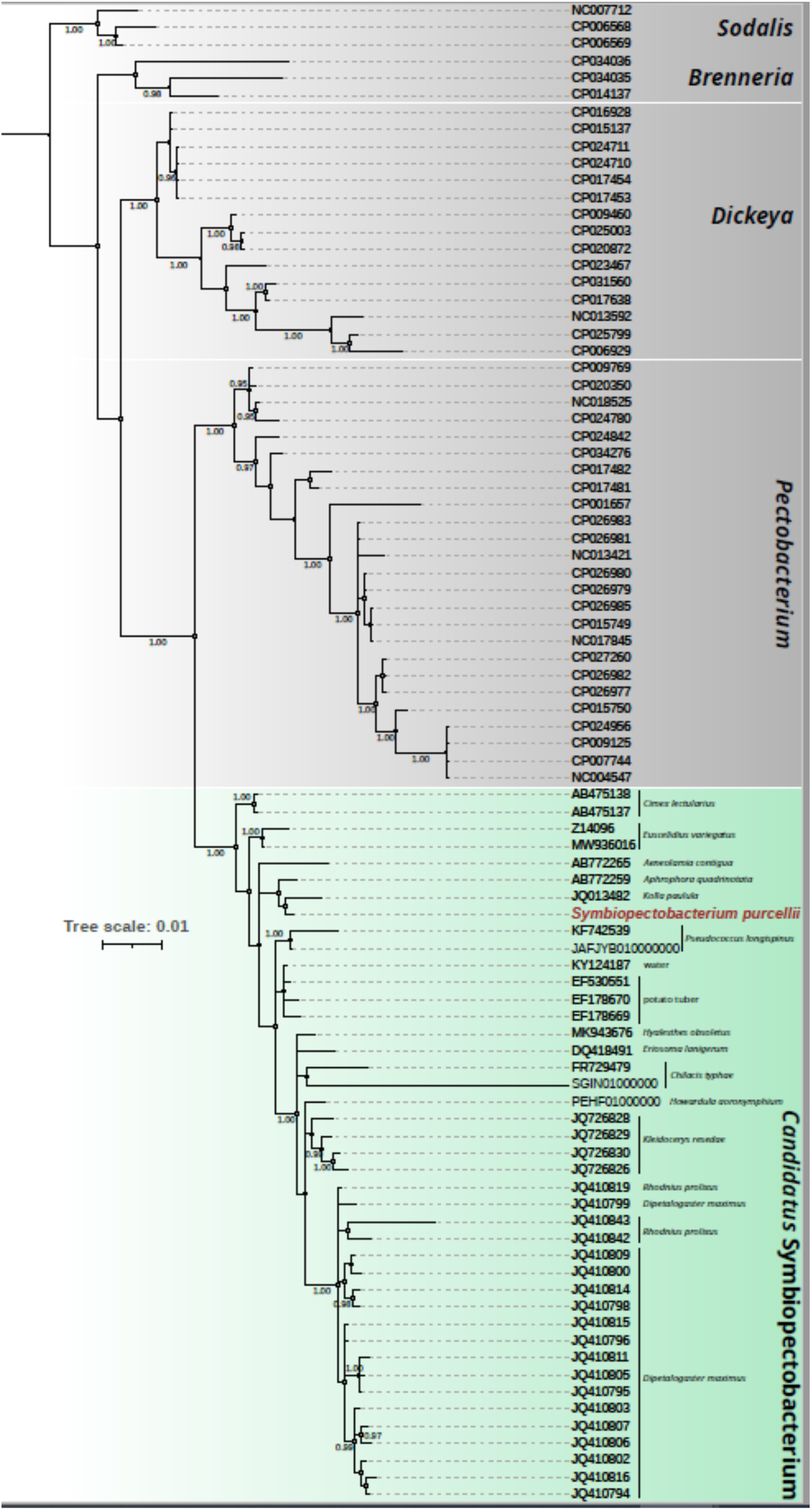
Phylogenetic affiliation of the 16S rRNA of *S. purcellii* compared to other strains, as estimated with Mr Bayes. Numbers on nodes represent posterior probability.

### In vitro Growth requirements

*S. purcellii* SyEd1 and *P. carotovorum* LMG 2404^T^ were grown at 30°C in BIOLOG GEN III plates. For *P. carotovorum* LMG 2404^T^, the presence of the purple tetrazolium dye as a result of growth and respiration in the wells where the strains had grown was visible after 24 h, while the wells that did not supported the growth of this strain remained colourless. In the case of *S. purcelliii* SyEd1, the plates had to be incubated for a total of 72h to allow bacterial growth. This is not surprising since, in our hands, *S. purcellii* SyEd1 grows slowly in BHI media (requiring up to 48h), and even slower in less rich media. Analyses indicated *S. purcellii* was considerably more fastidious than the comparator outgroup strain *P. carotovorum* LMG 2404^T^ in terms of metabolites that supported growth (Table 1) but had broader resistance to xenobiotics than this strain (Table 2). Growth conditions for *S. purcellii* on Biolog analysis was only modestly altered by addition of PGA.

**Table 1:**
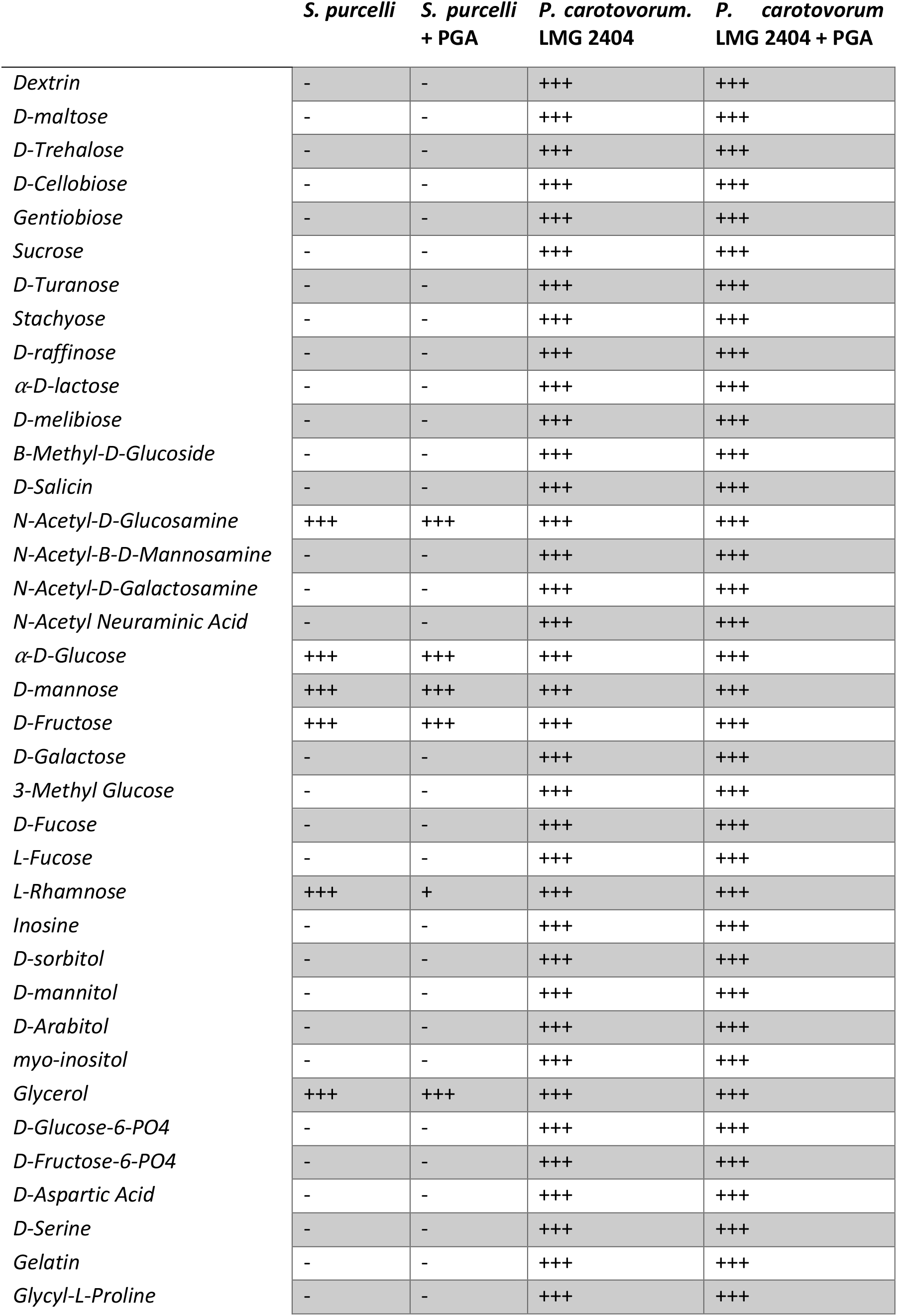
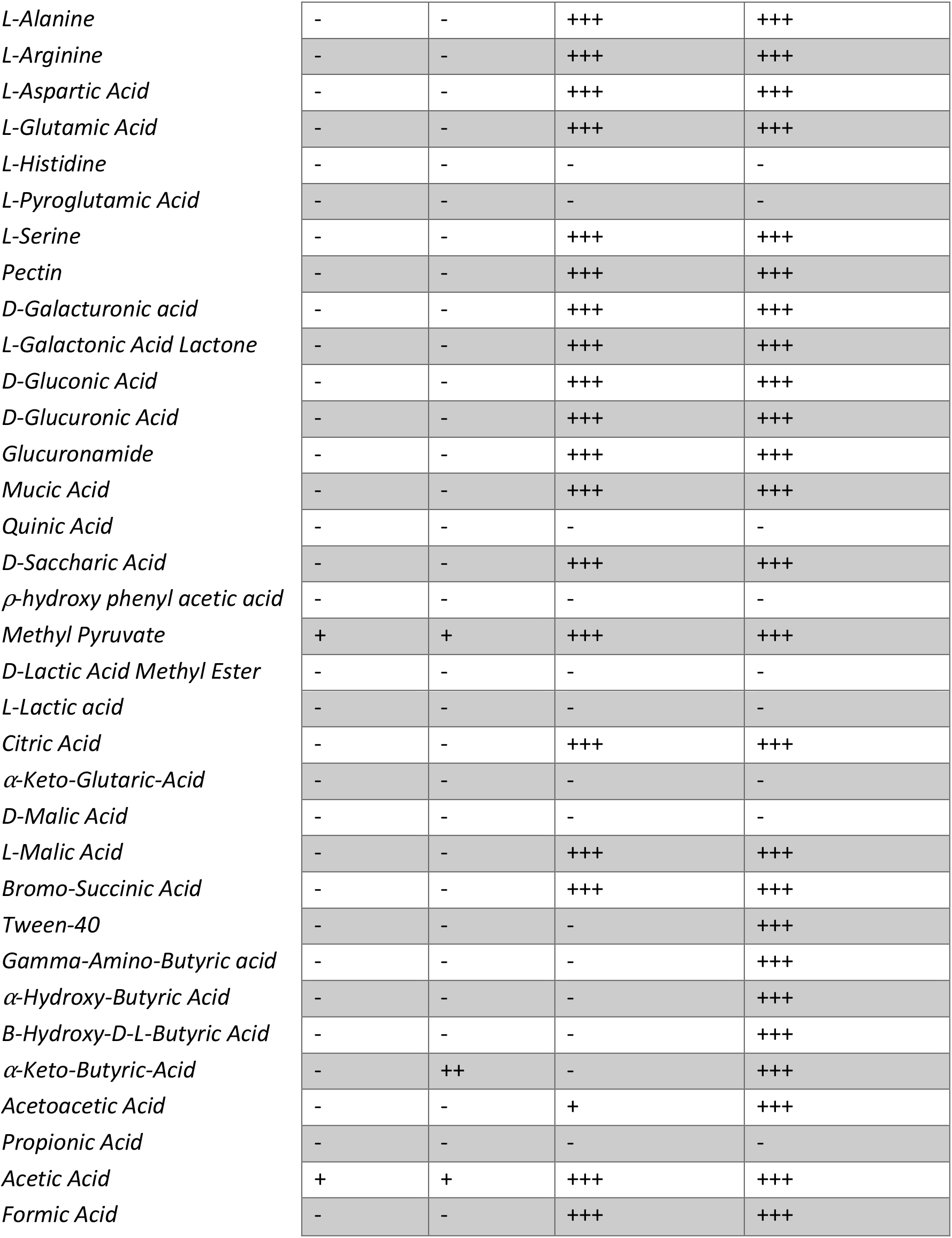
Utilization of carbon sources for Growth by *S. purcellii* in the presence and absence of PGA, with comparison to *Pectobacterium carotovorum* LMG 2404^T^. +++: strong growth; ++: medium growth; +: weak growth; and −: no growth.

**Table 2:** Impact of environmental and xenobiotic stress conditions on *S. purcellii* growth on Biolog III plates compared to *P. carotovorum,* in the presence/absence of PGA. +++: maintains full growth under condition stated, ++: medium growth; +: weak growth; and − :no growth under condition stated.

|  | *S. purcellii* | *S. purcellii*<br>+ PGA | *P. carotovorum*<br>LMG 2404<sup>T</sup> | *P. carotovorum*<br>LMG 2404<sup>T</sup> + PGA |
| --- | --- | --- | --- | --- |
| *pH 6* | +++ | +++ | +++ | +++ |
| *pH 5* | - | - | - | - |
| *1% NaCl* | +++ | +++ | +++ | +++ |
| *4% NaCl* | + | + | +++ | +++ |
| *8% NaCl* | - | - | - | - |
| *1% Sodium lactate* | +++ | +++ | +++ | +++ |
| *Fusidic acid* | +++ | +++ | +++ | +++ |
| *D-Serine* | - | +++ | - | - |
| *Troleandomycin* | +++ | +++ | +++ | +++ |
| *Rifamycin SV* | +++ | +++ | +++ | +++ |
| *Minocycline* | - | +++ | - | - |
| *Lincomycin* | ++ | ++ | +++ | +++ |
| *Guanidine HCl* | ++ | ++ | +++ | +++ |
| *Niaproof 4* | ++ | ++ | +++ | +++ |
| *Vancomycin* | +++ | +++ | +++ | +++ |
| *Tetrazolium Violet* | ++ | ++ | +++ | +++ |
| *Tetrazolium Blue* | +++ | +++ | +++ | +++ |
| *Nalidixic Acid* | ++ | ++ | - | - |
| *Lithium Chloride* | +++ | +++ | +++ | +++ |
| *Potassium Tellurite* | +++ | +++ | - | - |
| *Aztreonam* | +++ | +++ | - | - |
| *Sodium Butyrate* | ++ | ++ | +++ | +++ |
| *Sodium Bromate* | - | - | - | - |

### Potato infection assays

Potato slices infected with *S. purcellii* SyEd1 exhibited a complete absence of infection/ tissue maceration at the different time points tested (72h time point shown in Figure 3) in contrast to *P. carotovorum* LMG 02404^T^, used as positive control for infection. The assay was maintained for a week without any sign of infection being visible in *S. purcellii* SyEd1 infected potatoes.

**Figure 3:**
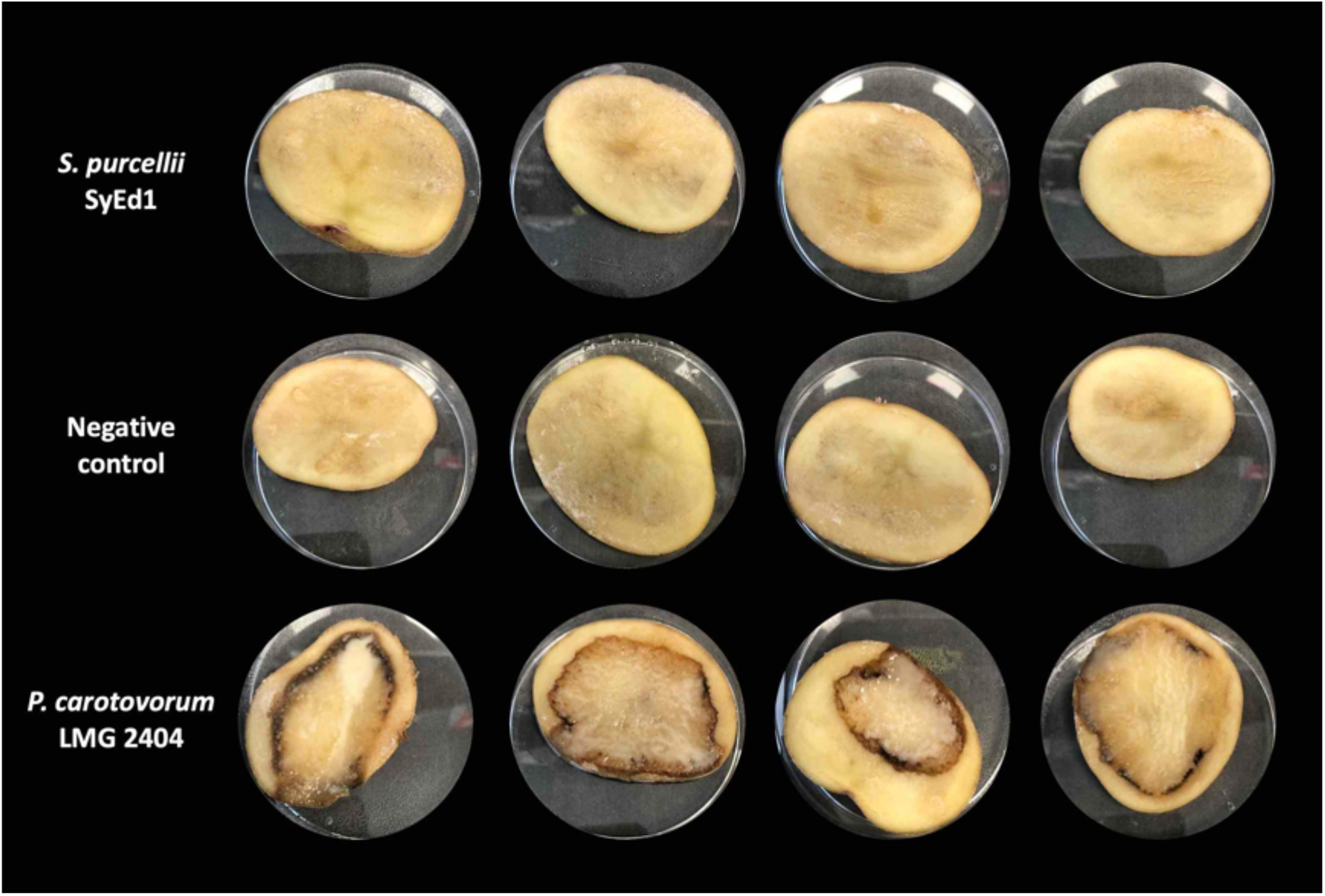
Virulence assay in potatoes.

### Genome sequence and assembly

The genome of the symbiont presented as a single circular chromosome of circa 4.9 MB with an average GC content of 52.5% (Table 3). No plasmids were identified. The complete predicted gene set consists of 4,494 protein-coding genes (including 312 predicted pseudogenes), 7 ribosomal RNA operons (5S, 16S, 23S) and 76 tRNAs. The average length of the protein-coding genes is 948 bp accounting for a coding density of about 86.2%. Pseudogenization rates were estimated to be circa 7% (312 predicted pseudogenes). The main chromosome was predicted to contain six intact prophage regions and three additional incomplete fragments. The complete genome assembly and the raw reads have been submitted to the DDBJ/EMBL/GenBank database under the BioProject accession number PRJNA756769 (genome accession number CP081864).

**Table 3.**
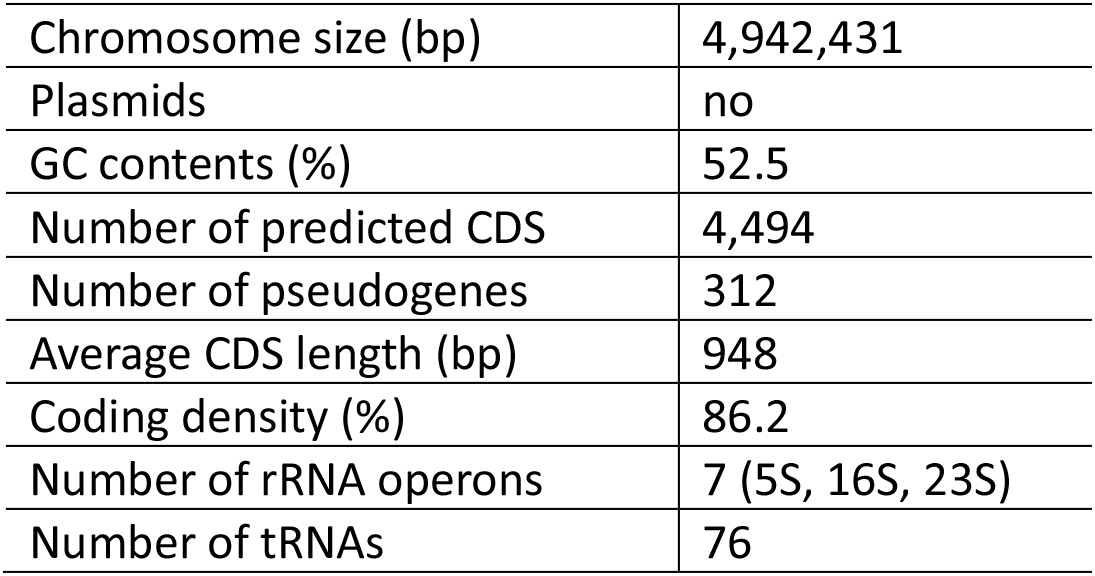
Genome features of the *Symbiopectobacterium purcellii* strain SyEd1 isolated from the leafhopper *Empoasca decipiens*.

|  |  |
| --- | --- |
| Chromosome size (bp) | 4,942,431 |
| Plasmids | no |
| GC contents (%) | 52.5 |
| Number of predicted CDS | 4,494 |
| Number of pseudogenes | 312 |
| Average CDS length (bp) | 948 |
| Coding density (%) | 86.2 |
| Number of rRNA operons | 7 (5S, 16S, 23S) |
| Number of tRNAs | 76 |

### Phylogenomic and functional analysis

To confirm the phylogenetic position of the *E. decipiens* symbiont we conducted a phylogenomic analysis base on the concatenated set of 527 single copy orthologue proteins across 56 related genomes (Figure 4). These results further support the placement of the symbiont in the *Cand.* Symbiopectobacterium clade.

**Figure 4:**
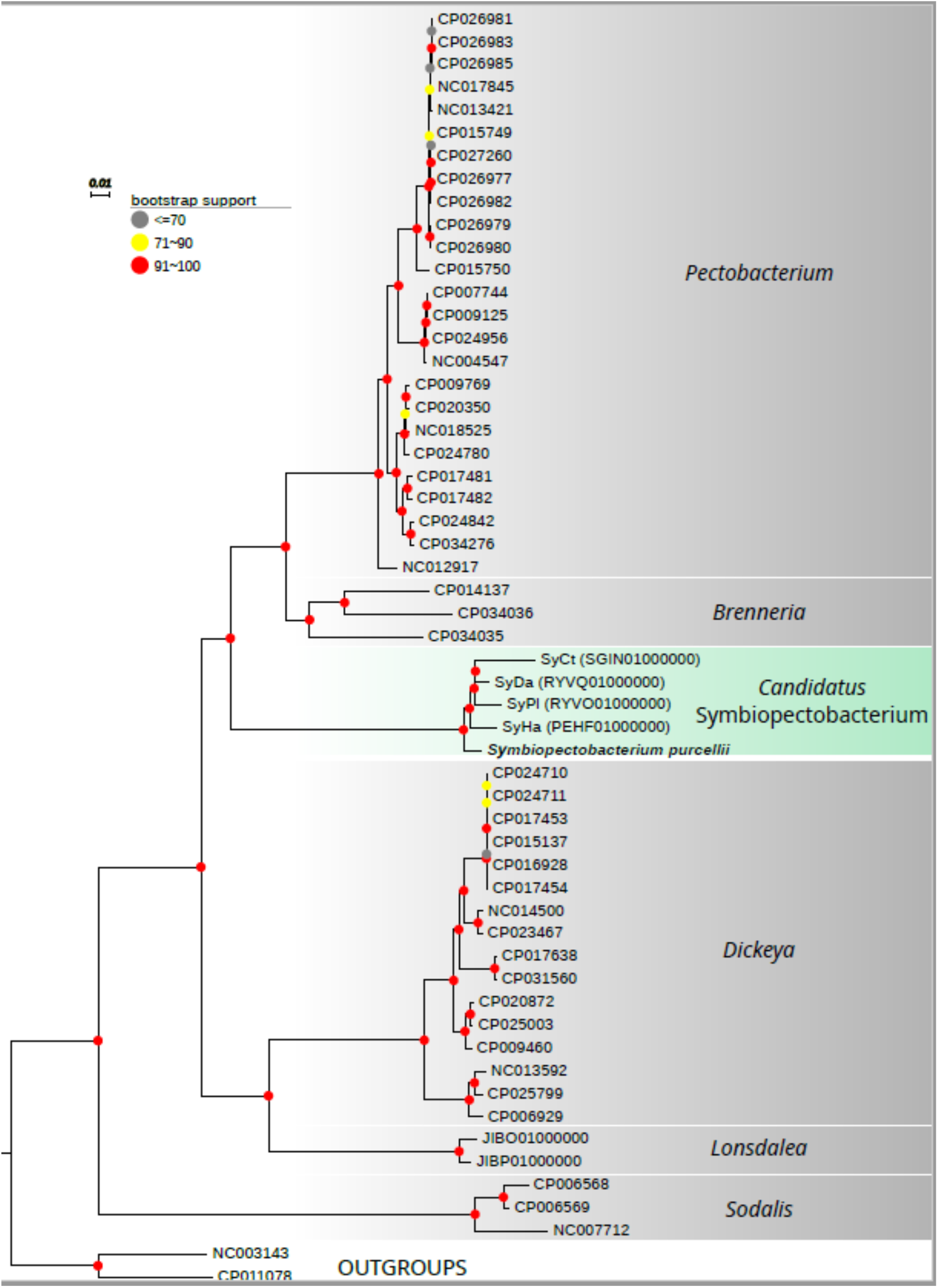
Affiliation of *S. purcellii* with other bacteria as estimated using IQTREE based on 527 shared single copy orthologues. Coloured dots on nodes represent bootstrap support.

The genome is predicted to encode type II, III and VI secretion systems alongside a wide array of predicted secreted toxins, compatible with its likely status as a symbiont of its insect host. Anti-SMASH predicted five genomic regions associated with small molecule production. Notable amongst these is a non-ribosomal peptide synthase (NRPS) region. It is unclear if the NRPS system produces siderophore molecules that permit growth in iron poor host environments, or antimicrobial compounds. In addition, anti-SMASH predicted putative gene clusters for the biosynthesis of thiopeptide, an aryl-polyene potentially providing defence against ROS, and betalactone synthesis. There is also a predicted homoserine lactone synthesis island *expI*/*expR1* that may be involved in sensing of microbial titre (see below); the genome encodes additional conserved elements of the Quorum sensing system, *expR2*, *gacA (expA)*, *gacS (expS)*, *rsmA* and *rsmB,* and *kdgR*. Finally, the genome encodes complete biosynthetic pathways for several B vitamins including thiamine (B1), riboflavin (B2), pantothenate (B5), biotin (B7), pyridoxine (B6) and folate (B9). A broad array of complete amino acid biosynthesis pathways was also observed, including serine, threonine, cysteine, methionine, valine, leucine, isoleucine, arginine, ornithine, arginine, proline, histidine, tryptophan, phenylalanine and tyrosine. Vitamin and amino acid provision are common means through which symbionts contribute to host function. There are also 5 *pel* genes predicted to encoded pectate lyase enzymes. The failure of the strain to utilize pectin on Biolog plates or on potato tubers may thus be context dependent.

### Analysis of homoserine lactones

*S. purcellii* SyEd1 analysis using LC-MS/MS revealed the presence of a single AHL that was characterised as *N*-(3-hydroxydecanoyl)-L-homoserine lactone (3-OH-C10-HSL). Figure 5 shows the LC-MS/MS chromatogram obtained from the *S. purcellii* SyEd1 sample compared to the 3-OH-C10-HSL standard and the uncultured BHI medium. Members of the genus *Pectobacterium* have been reported to produce 3-oxo-C6-HSL, 3-oxo-C8-HSL, C10-HSL (29), but, to the best of our knowledge, the presence of 3-OH-C10-HSL as the sole AHL in *S. purcellii* is unreported in related genera. This trait may help to identify novel members of this genus that we presume may have been previously misidentified as *Pectobacterium* spp. associated to the plants where the leafhoppers feed.

**Figure 5:**
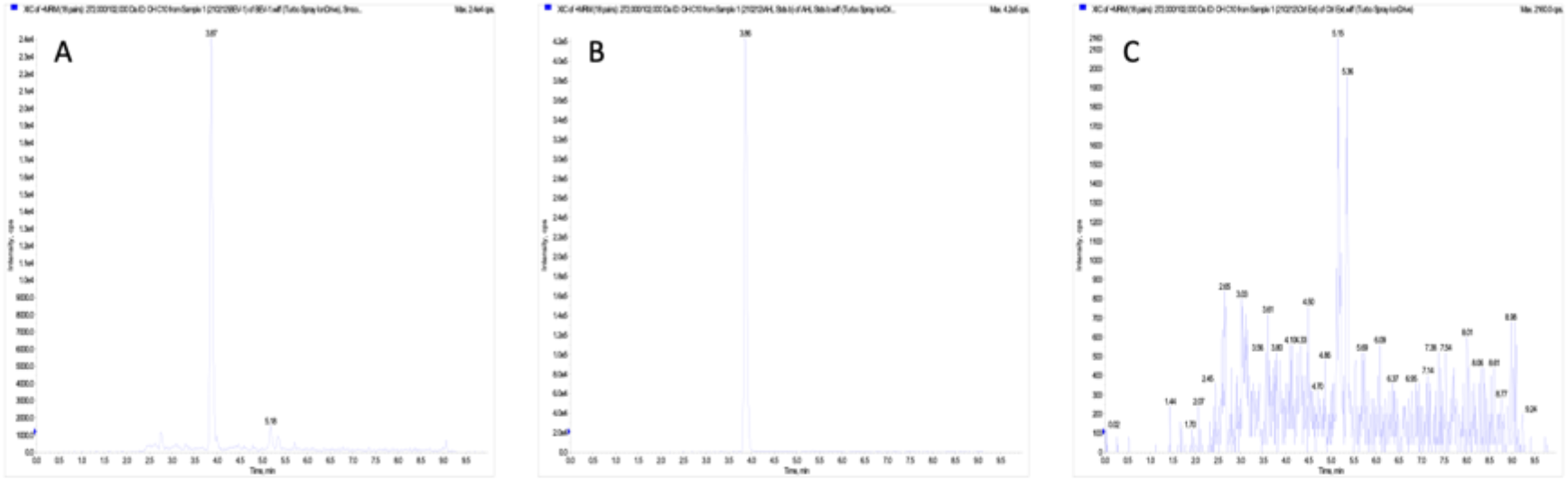
LC-MS/MS of an extraction of *S. purcellii* SyEd1 grown in BHI medium (A), 3-OH-C10-HSL standard (B) and negative control using a sterile BHI medium extract (C).

### Prevalence of *S. purcellii* in *E. decipiens* samples

Seven new *E. decipiens* specimens were collected and their identity confirmed by CO-1 amplification and sequencing. All samples were confirmed to be *E. decipiens* with >99.6% identity with previously deposited sequences in NCBI. The same gDNA extract was used to screen for the presence of the bacterial symbiont by PCR using our BEV_gyrB_F2 and BEV_gyrB_R2 primers. All seven samples produced an amplicon for *S. purcellii* (10), and the identity of the amplicon was confirmed through sequencing accounting for a 100% prevalence in the population tested (95%CI: 64% - 100%).

### Description of *Symbiopectobacterium purcelli gen. nov., sp. nov*

*Symbiopectobacterium purcellii*. *Symbiopectobacterium* (L. n. *sym bio pecto bacterium*) references the related *Cand.* Symbiopectobacterium that is an obligate symbiont of nematode worms, this name reflecting the symbiotic habit of the microbe, and the relationship of the genus as sister to *Pectobacterium*. The species name *purcellii* [pur.cell ii. L. m. gen.] is given in reference to Alexander Purcell, who isolated the first member of this clade, which he named the BEV symbiont (bacterium from *Euscelidius variegatus*).

Gram-negative rod-shaped bacterium. Grows optimally at 30 °C in BHI medium in the dark forming colonies within 24-48 h. Using Biolog GENIII plates, *S. purcelli* responded positively to the following carbon sources: D-glucose, D-mannose, D-fructose, glycerol, N-acetyl glucosamine, L-rhamnose, and weaker to methyl pyruvate and acetic acid. Growth was inhibited at pH5, by 4% and 8% NaCl, by D-serine, minocycline and sodium bromate. Growth was not impaired by 1% sodium lactate, fusidic acid, Troleandomycin, Rifamycin S, Lincomycin, Guanidine HCl, Vancomycin, Tetrazolium Violet, Tetrazolium blue, Potassium tellurite, Nalidixic Acid, Lithium Chloride, Aztreonam, Sodium Butyrate. The microbe does not cause macerations on potato slices.

*Symbiopectobacterium purcellii* gen. nov. sp. nov. form a cluster with a variety of uncultured symbionts of insects and nematodes, as well as the previously cultured strain BEV.

The type strain is SyEd1 (LMG 32449, CECT 30436) and was isolated from *Empoasca decipiens* (Hexapoda: Hemiptera: Cicadellidae) from Liverpool UK (53.3868° N, 2.9565° W). The genome consists of a single circular chromosome of size 4.9MB and DNA G+C content is 52.5 mol%. The 16S rRNA sequence of the type strain is available at accession OK044380. The complete genome assembly and the raw reads have been submitted to the DDBJ/EMBL/GenBank database under the BioProject accession number PRJNA756769 (genome accession number CP081864).

#### Acknowledgements

We are thankful to Alison Beckett (University of Liverpool, UK) for SEM Electron microscopy imaging service; Sam Edwards (University of Copenhagen, DK) for assistance in *E. decipiens* collection; Nigel Gotts (University of Liverpool, UK) for assistance evaporating the AHL-containing extracts; and Rita Valente (Instituto Gulbenkian de Ciência, PT) for useful discussions about the *Pectobacterium* virulence assays. This work was funded by a BBSRC grant to GH (grant BB/S017534/1). Miguel Cámara is partly funded by the National Biofilms Innovation Centre (NBIC) which is an Innovation and Knowledge Centre funded by the Biotechnology and Biological Sciences Research Council, Innovate UK and Hartree Centre (Award Number BB/R012415/1).

#### Conflicts of interest

The authors declare that there are no conflicts of interest.

## Notes

### Competing Interest Statement

The authors have declared no competing interest.

### Summary of Updates

Bacterial endosymbionts are found in multiple arthropod species, where they play crucial roles as nutritional symbionts, defensive symbionts or reproductive parasites. Recent work has highlighted a new clade of heritable microbes within the gammaproteobacteria that enter into both obligate and facultative symbioses, with an obligately required unculturable symbiont recently given the name Cand. Symbiopectobacterium. In this study, we describe a culturable rod shaped non- flagellated bacterial symbiont from this clade isolated from the leafhopper Empoasca decipiens. The symbiont is related to the transovarially-transmitted 'BEV' bacterium that was first isolated from the leafhopper Euscelidius variegatus by Alexander Purcell, and we therefore name the symbiont Symbiopectobacterium purcellii sp. nov. gen. nov. We further report the closed genome sequence for S. purcellii. The genome is atypical for a heritable microbe, being large in size, without profound AT bias and with little evidence of pseudogenization. The genome is predicted to encode Type II, III and VI secretion systems and associated effectors and a non-ribosomal peptide synthase array likely to produce bioactive small molecules. Predicted metabolism is more complete than for other symbionts in the Symbiopectobacterium clade, and the microbe is predicted to synthesize a range of B vitamins. However, Biolog plate analysis indicate metabolism is depauperate compared to the sister clade, represented by Pectobacterium carotovorum. A quorum-sensing pathway related to that of Pectobacterium spp. (containing an overlapping expI-expR1 pair in opposite directions and a "solo" expR2) is evidenced, and LC-MS/MS analysis reveals the presence of 3-hydroxy-C10-HSL as the sole N-acylhomoserine lactone (AHL) in our strain. This AHL profile is profoundly divergent from that of other Erwinia and Pectobacterium spp., which produce mostly 3-oxo-C6- and 3-oxo-C8-HSL and could aid group identification. Thus, this microbe denotes one that has lost certain pathways associated with a saprophytic lifestyle but represents an important baseline against which to compare other members of the genus Symbiopectobacterium that show more profound integration into host biology.

